# TCGA-Assembler 2: Software Pipeline for Retrieval and Processing of TCGA/CPTAC Data

**DOI:** 10.1101/214320

**Authors:** Lin Wei, Zhilin Jin, Shengjie Yang, Yanxun Xu, Yitan Zhu, Yuan Ji

**Affiliations:** Program of Computational Genomics & Medicine, NorthShore University HealthSystem, Evanston, Illinois, USA; Department of Applied Mathematics & Statistics, Johns Hopkins University, Baltimore, Maryland, USA; Department of Public Health Sciences, University of Chicago, Chicago, Illinois, USA

## Abstract

**Motivation:** The Cancer Genome Atlas (TCGA) program has produced huge amounts of cancer genomics data providing unprecedented opportunities for research. In 2014, we developed TCGA-Assembler (Zhu et al, 2014), a software pipeline for retrieval and processing of public TCGA data. In 2016, TCGA data were transferred from the TCGA data portal to the Genomic Data Commons (GDC), which is supported by a different set of data storage and retrieval mechanisms. In addition, new proteomics data of TCGA samples have been generated by the Clinical Proteomic Tumor Analysis Consortium (CPTAC) program, which were not available for downloading through TCGA-Assembler. It is desirable to acquire and integrate data from both GDC and CPTAC.

**Results:** We develop TCGA-Assembler 2 (TA2) to automatically download and integrate data from GDC and CPTAC. We make substantial improvement on the functionality of TA2 to enhance user experience and software performance. TA2 together with its previous version have helped more than 2,000 researchers from 64 countries to access and utilize TCGA and CPTAC data in their research. Availability of TA2 will continue to allow existing and new users to conduct reproducible research based on TCGA and CPTAC data.

**Availability:** http://www.compgenome.org/TCGA-Assembler/

## 1 Introduction

Over the past decade, The Cancer Genome Atlas (TCGA) program has generated large amounts of genomics and proteomics data characterizing the molecular landscape of more than 11,000 tumors across 33 cancer types. TCGA data provide unprecedented opportunities for systematic investigation of cancer mechanisms at multiple molecular and regulatory layers and has been used in numerous cancer studies (Tomczak et al, 2015; Cancer Genome Atlas Network, 2012).

In 2014, we developed TCGA-Assembler (TA), an open-source R software pipeline for retrieving, assembling and processing public TCGA data (Zhu et al, 2014). TA was developed to communicate with the TCGA Data Portal, the official repository of TCGA data at that time. In July 2016, TCGA Data Portal was terminated and all TCGA data were transferred to the newly established Genomic Data Commons (GDC, https://gdc.cancer.gov/), which made TA impossible to function for its users. In addition, new proteomics data of TCGA samples have been recently generated and released by the Clinical Proteomic Tumor Analysis Consortium (CPTAC) (Zhang et al, 2016), a different data repository. The CPTAC program utilizes the mass spectrometry proteomics technology of isobaric Tags for Relative and Absolute Quantification (iTRAQ) (Ross et al, 2004) to generate whole-proteome measurements. Thus, it is desirable to revive the functions of TA for its existing and future users and integrate CPTAC data with TCGA data.

To this end, we develop TCGA-Assembler 2 (TA2) for the cancer research community. Like TA, TA2 possesses two software modules. Module A is completely rebuilt using the GDC application programming interfaces (Fig. 1). Compared to TA, Module A in TA2 adds new functions that allow download of TCGA somatic mutation data and proteomics data from CPTAC. Module B has been enhanced with new functions and features, such as preprocessing of somatic mutation and proteomics data. In TA 2 we have also optimized existing functions to improve their performance, such as implementing the batch downloading mechanism to reduce the chance of data transferring error over Internet.

**Figure 1.**
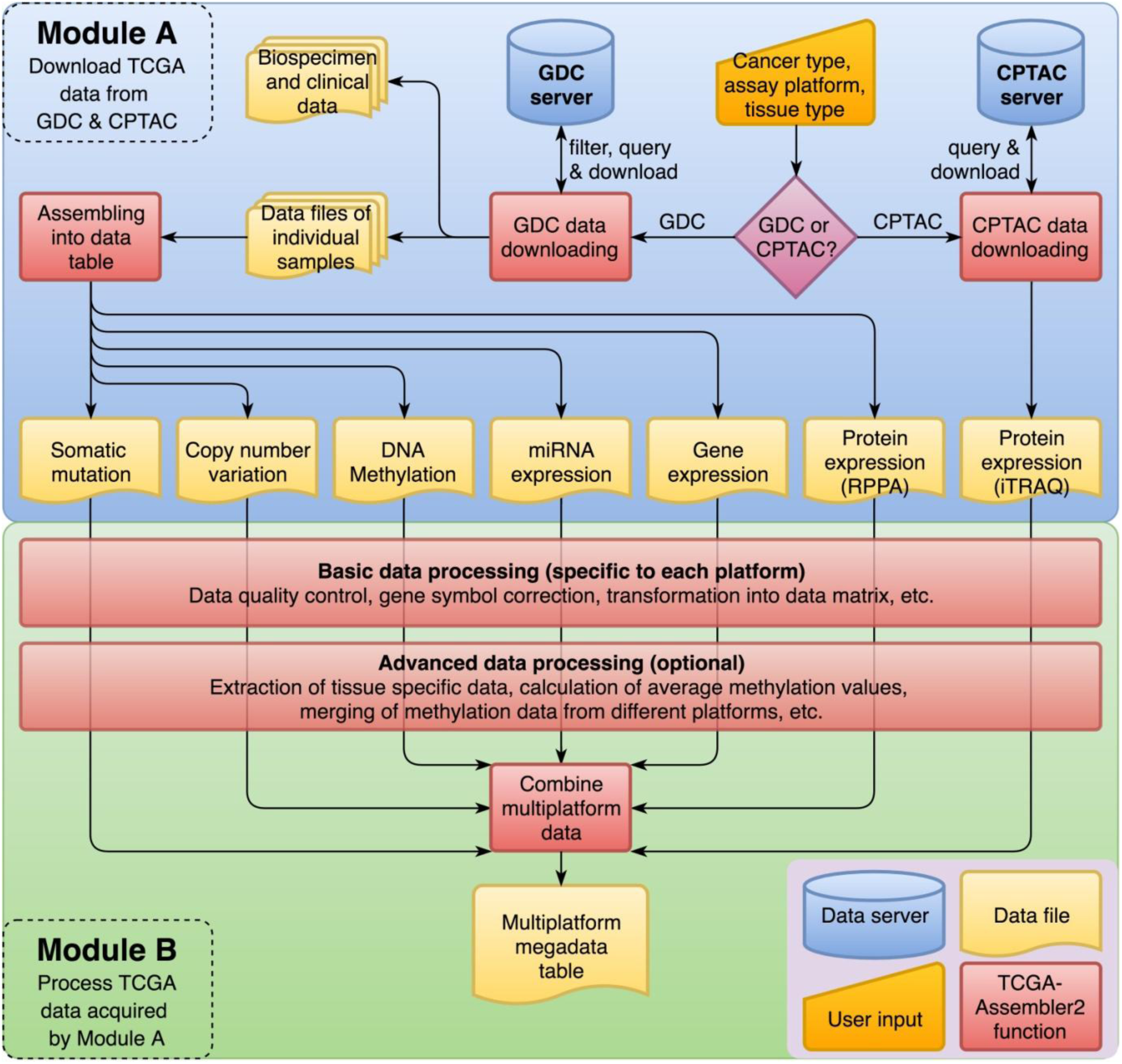
Software architecture and functional flowchart of TA2.

## 2 Methods

### Module A

Module A acquires data files according to user's specification, and provides one function for downloading each data modality, including gene expression, miRNA expression, DNA copy number, DNA methylation, somatic mutation, RPPA (Reverse Phase Protein Array) protein expression, iTRAQ protein expression (from CPTAC), and clinical and biospecimen information (Fig. 1). Through input arguments, user can specify the cancer type of interest (such as lung cancer), and optionally the assay platform used to generate the data, the tissue type of interest (such as primary solid tumor or blood derived normal), and the TCGA barcodes of patients. Based on the input TA2 filters the data files and downloads the requested data from the GDC and CPTAC data severs. In addition, Module A will integrate the data files belonging to the same cancer type and assay platform into a single data table. Section 1 of the Supplementary Information gives details about how these data downloading and assembly functions are implemented.

### Module B

Module B includes functions for processing the data acquired by Module A to fulfill various data processing needs (Fig. 1). Module B functions are divided into two groups: basic data processing functions and advanced data processing functions. For each of the data modalities, the basic processing function performs quality control on data (such as validate/correct gene symbols and produce data visualization for identifying outliers), removes redundant genomic feature descriptions, ensures that each row in a data table corresponds to a unique genomic feature, and separates different types of measurements into their own data tables. Besides these functionalities, basic processing functions of specific data platforms can summarize data into measurements suitable for subsequent analysis, such as the gene-level copy number alternation and the count of somatic mutations in a gene across loci. The advanced data processing functions perform further data manipulation and calculation based on data files produced by the basic data processing functions (Fig. 1), such as calculating the average DNA methylation value of CpG sites in a particular region of gene (e.g. the promoter region) and integrating multiple data platforms of samples into a mega data table (Table S1), where the multi-platform data of a gene are organized as adjacent rows in the table. Such a mega data table is particularly useful for cross-platform, integrative analysis of TCGA data (Zhu *et al,* 2015), which is one of the primary goals of TCGA program. Section 2 of the Supplementary Information gives more details about Module B functions.

## 3 Results

Table S2 summaries the number of patient samples for which data can be retrieved and processed using TA2. Compared to the original TA, new data modalities that are available through TA2 include the TCGA somatic mutation data and the iTRAQ proteomics data from CPTAC that include proteome, phosphoproteome, and glycoproteome measurements. Currently, CPTAC only has data for four TCGA cancer types, including breast invasive carcinoma, colon adenocarcinoma, ovarian serous cystadenocarcinoma, and rectum adenocarcinoma, and is expected to cover more TCGA cancer types in the future. Besides providing more data, TA2 also modified and optimized functions to improve their performance and reliability. For example, for cancer types with a lot of samples and assay platforms with huge amounts of features, TA2 exploits batch downloading, in which the target samples are divided into batches and downloaded sequentially. This significantly reduces the chance of getting a data transferring error compared to transferring a single large data file over Internet. See Sections 3 and 4 in the Supplementary Information for more information of TA2 software package, its future development plan, and its relationship to other resources and tools.

## Acknowledgements

We thank Dr. Robert Grossman and GDC team for providing discussion and comments during the development of TA2.

## Funding

The research is partly supported by NIH 2R01 CA132897.

## Conflict of Interest

none declared.

